# Mouse pup ultrasonic vocalization repertoires shift with age and behavioral context across the first postnatal week

**DOI:** 10.64898/2026.11.24.768928

**Authors:** Jacob Elrod, James Buchanan, Kevin J. Reilly, Devin Casenhiser, Billy You Bun Lau, Keerthi Krishnan

## Abstract

Pup ultrasonic vocalizations (USVs) are thought to communicate and elicit maternal care, yet the structure of the pup call repertoire and how it changes across the vulnerable first postnatal week remain poorly characterized. Here we analyzed approximately 59,000 USVs emitted by wildtype mouse pups (postnatal days P0–P5) during a pup retrieval paradigm that included habituation, pup isolation, and retrieval phases, performed with wild-type (WT) and *Mecp2*-heterozygous (Het) female mice as alloparental surrogates. Using DeepSqueak-based acoustic feature extraction and unsupervised clustering, we identified eight distinct call types, grouped into two broad classes based on shared acoustic properties. The relative prevalence and sequencing of these call types shifted systematically with postnatal age and behavioral phase, indicating that the pup vocal repertoire is both developmentally and contextually dynamic during the first postnatal week. However, call-type composition and acoustic characteristics did not differ with behavioral outcome, consistent with the model in which USVs function as general arousal signals integrated with other sensory cues. A candidate sequence-level pattern distinguished failed retrievals but was based on few calls and requires further validation. Pup call repertoire and developmental trajectory were also indistinguishable between WT and Het surrogates, indicating that the previously reported retrieval deficit in Het females does not arise from atypical pup vocal signals. Together, these results provide a detailed developmental and contextual map of the early pup call repertoire and a framework for relating call-type dynamics to auditory circuit plasticity across early postnatal life.

## Introduction

Rodent ultrasonic vocalizations (USVs) are thought to be important for communication across life stages and have been studied extensively to model social communication. Pups emit USVs predominantly when separated from the nest, and are crucial signals to elicit maternal care (D’Amato et al., 2005; Portfors & Perkel, 2014; Smotherman et al., 1974). Characterization of pup call types in this behavioral context is relatively understudied in the first postnatal week, when pups cannot thermoregulate and are dependent on maternal care for survival (Smotherman et al., 1974; Wöhr et al., 2008). Existing call classification efforts have predominantly relied on calls during isolation, and sampled at non-consecutive ages, rather than at daily resolution in ethologically relevant behavioral contexts (Castellucci et al., 2018; D’Amato et al., 2005; Wöhr et al., 2008). USV playback behavioral experiments and functional studies in the auditory cortex of rodent dams and sensitized females have shown that pup USVs elicit affiliative behaviors, likely mediated by specific intrinsic and synaptic plasticity in pyramidal and parvalbumin+ GABAergic interneurons (Cohen et al., 2011; Cohen & Mizrahi, 2015; D’Amato et al., 2005; Elwood & Keeling, 1982; Galindo-Leon et al., 2009; Lau et al., 2020a; Lin et al., 2013; Liu & Schreiner, 2007; Marlin et al., 2015; Screven & Dent, 2019; Shair et al., 1998; Tasaka et al., 2018, 2025). In most of these electrophysiological studies, a limited number of USVs are played back repeatedly as stimuli. These stimulus sets of about 8 calls are typically drawn from a narrow postnatal window, leaving it unclear whether these calls are representative of what the auditory system encounters during pup retrieval behavior. Furthermore, it is unknown if USV call characteristics change developmentally over this critical first week of life. These two open questions converge on a fundamental question: whether acoustic features of the pup USVs or the adult receiver’s perception and interpretation of those calls, form the primary basis for social communication in this first week of life. Additionally, many studies now use USVs as a readout for social communication in animal models of neurological disorders (Fujita et al., 2008; Moreno et al., 2023; Premoli et al., 2021; Screven & Dent, 2019; Yin et al., 2016). Most of the studies use USV counts at a particular age, typically in a pup isolation condition to elicit the most vocalizations, as the chief metric. Thus, behavioral context, which is key for USVs to be a social communication signal, is often lacking.

Here, we use the adult female *Mecp2-*heterozygous (Het) mouse model of Rett syndrome to test if acoustic features of pup USVs are important for efficient pup retrieval behavior. We have previously shown that Het mice exhibit inefficient retrieval behavior, altered trajectory patterns only in the retrieval phase (not in habituation or adult isolation phases), and atypical ethogram, while compared to wildtype (WT) littermate controls (Krishnan et al., 2017; Stevenson et al., 2021; Mykins et al., 2024). Furthermore, auditory cortex neurons of Het show atypical plasticity response with pup USV stimuli, and not broad band noise (Krishnan et al, 2017; Lau et al, 2020a). This result points to the adult receiver’s perception and interpretation to be a cause for the inefficient pup retrieval. However, it is also possible that Het mice fail to adapt appropriately to likely developmental changes in pup calls over days. Alternatively, if pups emit a “come get me” signal that drives retrieval, Het mice may fail to perceive or interpret that signal correctly. These distinctions are important as Rett syndrome patients exhibit issues in both sensory processing and vocal communication among other symptoms (Sysoeva et al., 2020).

To investigate this hypothesis, we analyzed ∼59,000 USVs emitted by wildtype pups across the pup retrieval paradigm performed by alloparental WT and Het for 6 days (pups aged P0-P5), using DeepSqueak (Coffey et al., 2019). We identified 8 call clusters that showed distinct developmental changes across days and behavioral contexts. However, pup USV call clusters and characteristics were largely similar regardless of behavioral outcome or surrogate genotype. Thus, we were unable to identify specific “come get me” calls. However, we did identify a candidate sequence-level pattern that distinguished failed retrievals in early postnatal days, but it was based on very few calls and requires further validation. These results indicate that the atypical retrieval behavior in Het is unlikely to arise from pup USVs they receive, thus strengthening the receiver-side deficits concept, as shown previously (Lau et al., 2020a). Most importantly, the dynamic developmental changes we identify in call cluster prevalence provides a tool for studying auditory cortex plasticity longitudinally.

## Methods

### Animals

Two mouse strains, CBA/CaJ (RRID:IMSR_JAX:000654) and *Mecp2^heterozygous^* (Het) (C57BL/6J, B6.129P2(C)-*Mecp2^tm1.1Bird^*/J, RRID:IMSR_JAX:003890) (Guy et al., 2001) were used in the study. All animals were group-housed by sex after weaning, raised on a 12/12-h light/dark cycle (lights on at 7 am), and received food and water ad libitum. All procedures were conducted in accordance with the National Institutes of Health’s Guide for the Care and Use of Laboratory Animals and approved by the Institutional Animal Care and Use Committee at the University of Tennessee-Knoxville.

### Pup Retrieval behavior

Pup retrieval behavior was performed as previously described (Stevenson et al., 2021). Briefly, one Het virgin female mouse and one *Mecp2^WT^* virgin female mouse (both referred to henceforth as *surrogates*), which create a cohort, were cohoused with pregnant CBA/CaJ female 3-5 days before birth. Behavioral experiments were performed using female Het and *Mecp2^WT^* siblings (WT) at 10–12 (adult) weeks of age between the hours of 9 am and 6 pm (during the light cycle). Nine cohorts of surrogates were put through the pup retrieval assay upon birth of the pups (postnatal day 0, P0). The assay is comprised of 4 phases: habituation (Hab), adult isolation, pup retrieval (Ret) and pup isolation (Iso). First, in habituation one surrogate was habituated with 3-5 pups in the nest of the home cage for 5 min; then in adult isolation the pups are removed, leaving the surrogate alone in the nest for 2 minutes; finally, one pup is placed in each corner and one in the center of the cage. If there were <5 pups, the nest would be left empty. The surrogates have a maximum of 5 minutes to complete the retrieval phase. While one of the females was performing the pup retrieval task, the mother, her pups, and the other adult were housed in a separate new cage. We also recorded scattered pups in the absence of adults, termed pup isolation, from P0-P5, for 5 minutes. All animals were returned to the home cage after testing. Trials were completed for 6 consecutive days (P0-P5). All behavioral assays were performed in the dark box and were recorded using a Foscam Wired IP Camera. USV audio was simultaneously recorded using 4 ultrasonic microphones (Avisoft; Glienicke, Germany) hung in each corner of the home cage during each phase.

### DeepSqueak

The DeepSqueak software package (Coffey et al., 2019) (version 3.1) was used to parse and analyze the USV recordings. The parsing of individual USVs was carried out using Mouse YOLO detector V3 model available in DeepSqueak. USV detections across all phases (except adult isolation) totaled ∼59000 calls. Surrogate WT and Het only emitted a total of 13 calls in isolation. The adult call structure was also distinct from pup USVs, and thus were not included in this analysis. USVs were then clustered via unsupervised clustering using contour invariant variational autoencoder (Goffinet & Pearson, 2021; Sainburg et al., 2020). All default settings were used for training autoencoder (250 training iterations). ELBO-loss method was utilized to determine the optimal number of clusters. This method yielded 8 clusters (setting: Max Clusters: 100, Replicate: 3). The model was validated using 80 percent of the data, as well as a “leave-on-out” validation method. The characteristics of the USV clusters were determined by deriving 7 call features: call length, principal frequency, low frequency, high frequency, delta frequency, slope, and sinuosity. The USV features and corresponding cluster labels were exported from DeepSqueak to R and RStudio (Racine, 2012) for analysis and visualization. Call rate for retrieval was calculated by dividing the number calls by the time it took surrogates to return all pups to the nest in successful trials, and by the full 300s in failure trials. Call rates for habituation and pup isolation were calculated by dividing number calls by 300 seconds. Cluster distance from center was generated from DeepSqueak’s built-in cluster distance analysis.

### Statistical Analysis

All data were processed and analyzed using R and RStudio (R Core Team, 2021) and figures generated in Adobe Illustrator.

To evaluate developmental changes in vocalization behavior (Figure 1), linear regression analyses were performed with postnatal day treated as the predictor variable and call count or call rate as the dependent variable. Analyses were performed separately for individual behavioral clusters and experimental phases to determine whether vocalization measures exhibited significant linear changes across developmental days. Developmental day was represented numerically according to the ordered postnatal sequence, with P0 through P5 corresponding to successive developmental time points.

**Figure 1.**
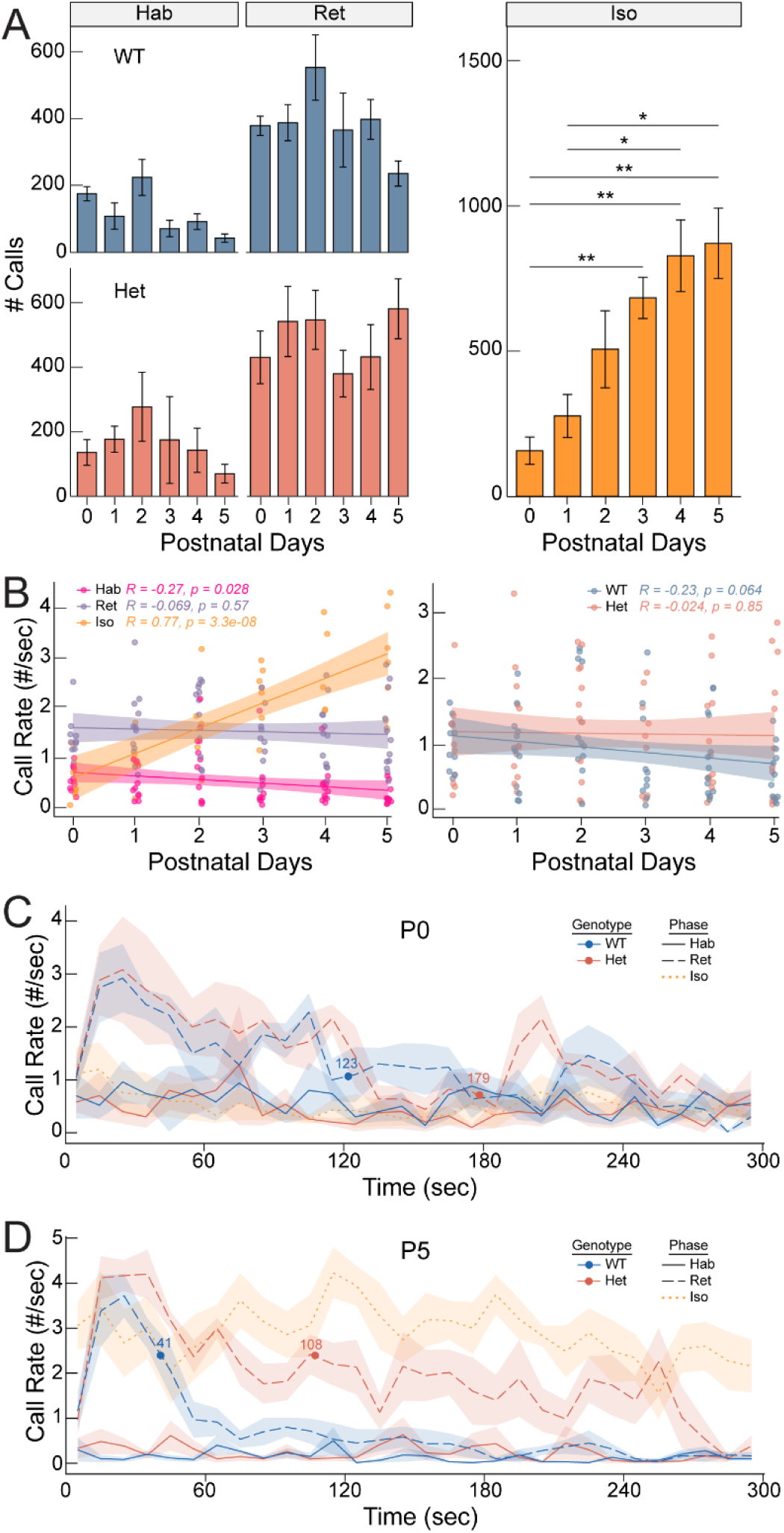
Phase and day-specific dynamics in pup calls across different contexts during pup retrieval task: **A)** Bar plots depicting call counts by postnatal day in habituation (Hab) and retrieval (Ret) for WT (blue), Het (red), and isolation (Iso, right, yellow). WT and Het showed no significant change between days in either phase. Isolation showed significant increases from P0 to P3, P4 and P5, as well as P1 to P4 and P5. *\*p<0.05, **p<0.001, FDR corrected Dunn’s test.* Bars represent mean ± S.E.M. **B)** Calls per second (rate) by days for each phase (left) and genotype (right). In phase comparisons, Hab (pink) showed consistently lower call rate across days of behavior compared to Ret (purple), which remained stable across days. Iso (yellow) significantly increased across days. In genotype comparisons, both WT and Het showed consistent call rate (combined Hab and Ret) across days that were not significantly different from each other. Lines plus shadow indicate linear regression ± 95% SEM. **C-D)** Line plots depicting calls binned per 10 seconds for WT (blue) and Het (red) in Hab (solid line) and Ret (dashed line) on P0 (C) and P5 (D) for the full 300 seconds of the phase, with circles indicating average end of retrievals for each genotype. Iso (yellow, dotted) calls are binned similarly. Pup calls during WT retrieval showed decreases across the behavior from P0 to P5, while call rates during Het retrieval increased across duration from P0 to P5, to almost isolation levels. Iso call rates increased from P0 to P5. Hab call rates remained low over days. Hab: n= 9,417, Ret: n = 29,837, Iso: n = 20,028.

For each regression, the fitted linear relationship between developmental day and the vocalization measure was used to estimate the direction and magnitude of the developmental trend. Positive regression coefficients indicated an increase in calling across developmental days, whereas negative coefficients indicated a decrease. The strength and statistical significance of the association were evaluated using the model correlation and associated p-value. Regression lines and their corresponding 95% confidence intervals were used for visualization of developmental trajectories.

Chi squared tests were used to compare cluster frequency between genotypes (Figure 2, 4). Categorical behavioral measures, including behavioral classifications and outcome categories, were summarized as frequencies and proportions. Differences in the frequencies of call clusters across phases and genotypes were assessed using Pearson’s chi-squared tests of independence. For significant chi-squared tests, standardized residuals were examined to identify specific categories contributing to the observed associations. Standardized residuals were used to evaluate deviations between observed and expected frequencies and to characterize the direction and magnitude of differences for individual comparisons.

**Figure 2A-C.**
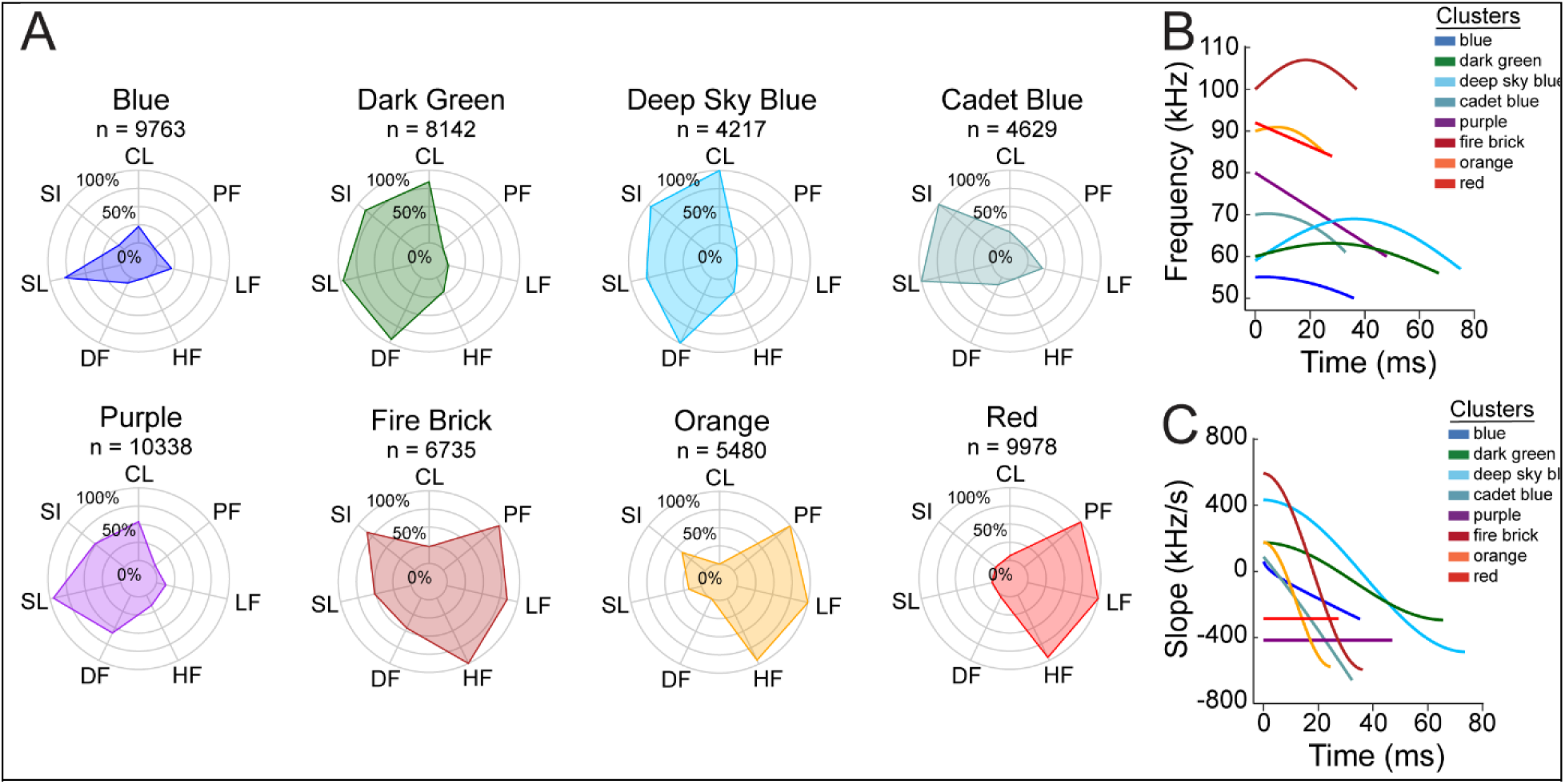
DeepSqueak call clusters are distinct in metrics, separating into ‘cool’ and ‘warm’ categories: **A)** Radar plots depicting average DeepSqueak metrics for each cluster scaled from 0-100, with number of calls for each cluster. Cool clusters (blue, purple, deep sky blue, cadet blue, dark green) share similar slope, sinuosity, and call length. Warm clusters (fire brick, yellow, red) share similar high frequency. CL = Call Length, PF = Principal Frequency, LF = Low Frequency, HF = High Frequency, DF = Delta Frequency, SL = Slope, SI = Sinuosity. **B)** Graph showing most representative call of each cluster, based on frequency and time. **C)** Graph plotting most representative call of each cluster based on slope over time.

To account for multiple statistical comparisons across phases, behavioral measures, and experimental groups, p-values were adjusted using the false discovery rate (FDR). FDR-adjusted p-values < 0.05 were considered statistically significant. All statistical tests were two-sided.

Spearman’s correlational analysis was performed to calculate correlational analysis between call clusters and latency score (metric described previously Stevenson et al, 2021) to determine if cluster impacts retrieval ability of surrogates (Figure 4). Correlations between call rate and latency score were assessed using Pearson correlation coefficients, computed separately for each postnatal day and genotype. To account for multiple comparisons across the day-by-genotype combinations, p-values were adjusted using Benjamini-Hochberg false discovery rate. The adjusted value of p < 0.05 was considered significant for each statistical test. Calls during the success or failed retrievals were used for calculating correlation with behavioral outcome and efficiency. To label successes and failures, events were timestamped from initial interaction with the pup (whisker length away) until pup was successfully retrieved or dropped in the case of failures. This time window was determined manually using timestamps.

**Figure 3:**
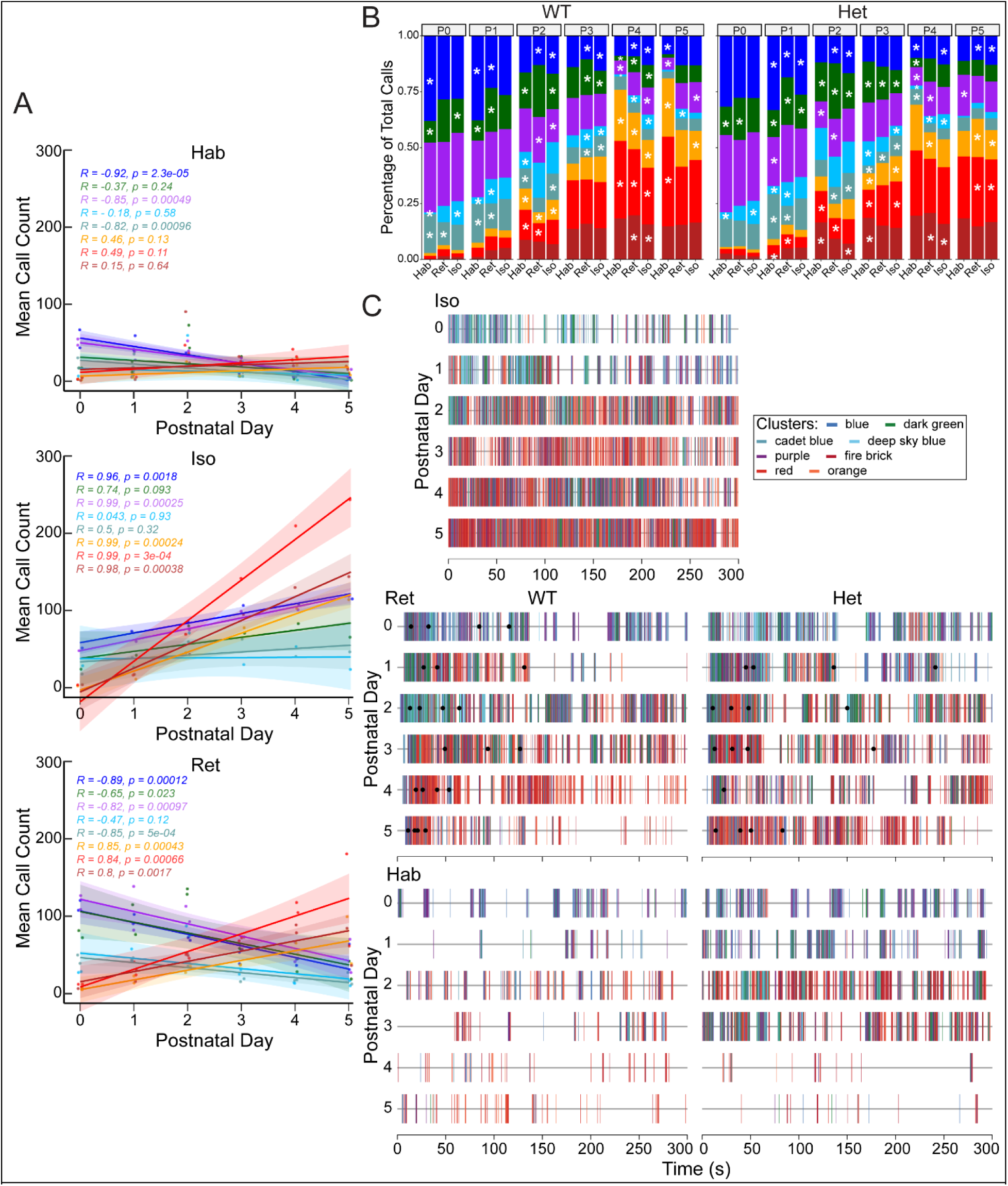
Call clusters are significantly changed across phases of behavior and days. **A) Top:** Linear regression of each cluster’s average calls across postnatal days P0-P5 in habituation phase. 5 out of 8 clusters showed no significant correlation across days. Only blue, purple and cadet blue showed significant negative correlation. **Middle:** Linear regressions of each cluster’s call rate in isolation phase. Warm clusters showed significant positive correlations across days, while most cool clusters showed significant positive correlation. Dark green (p=0.093) and cadet blue (p= 0.32) were the exceptions. Together, this pattern suggests an overall increase in number of calls for most clusters during isolation over days. **Bottom:** Linear regression of each cluster’s average calls in retrieval phase. All warm clusters showed significant positive correlation across days, while all cool clusters except deep sky blue showed significant negative correlation, indicating a decrease in mean call count across days. Deep sky blue showed no significant correlation. B) Bar plots depicting each cluster as a percentage of total calls in each genotype for all three phases across days. Significant differences in various clusters were shown between all phases across days (* denotes a phase’s cluster is significantly different from the expected frequency between all three phases on that day. *FDR corrected chi-squared test, * = p< 0.05*). Changes between call clusters indicate pups call differently in different contexts, but developmental trajectory remains similar. C) Sequence plots of representative cohort showing individual calls across phases (top: iso; middle: ret; bottom: hab). All phases showed developmental trend of cool to warm color clusters over days. Density of calls was highest in iso. Number of calls typically decreased after retrieval in both WT and Het. Calls are colored by cluster. Ret: black dots indicate the start of a successful retrieval, and green dots indicate the last successful retrieval on a day.

**Figure 4A-B.**
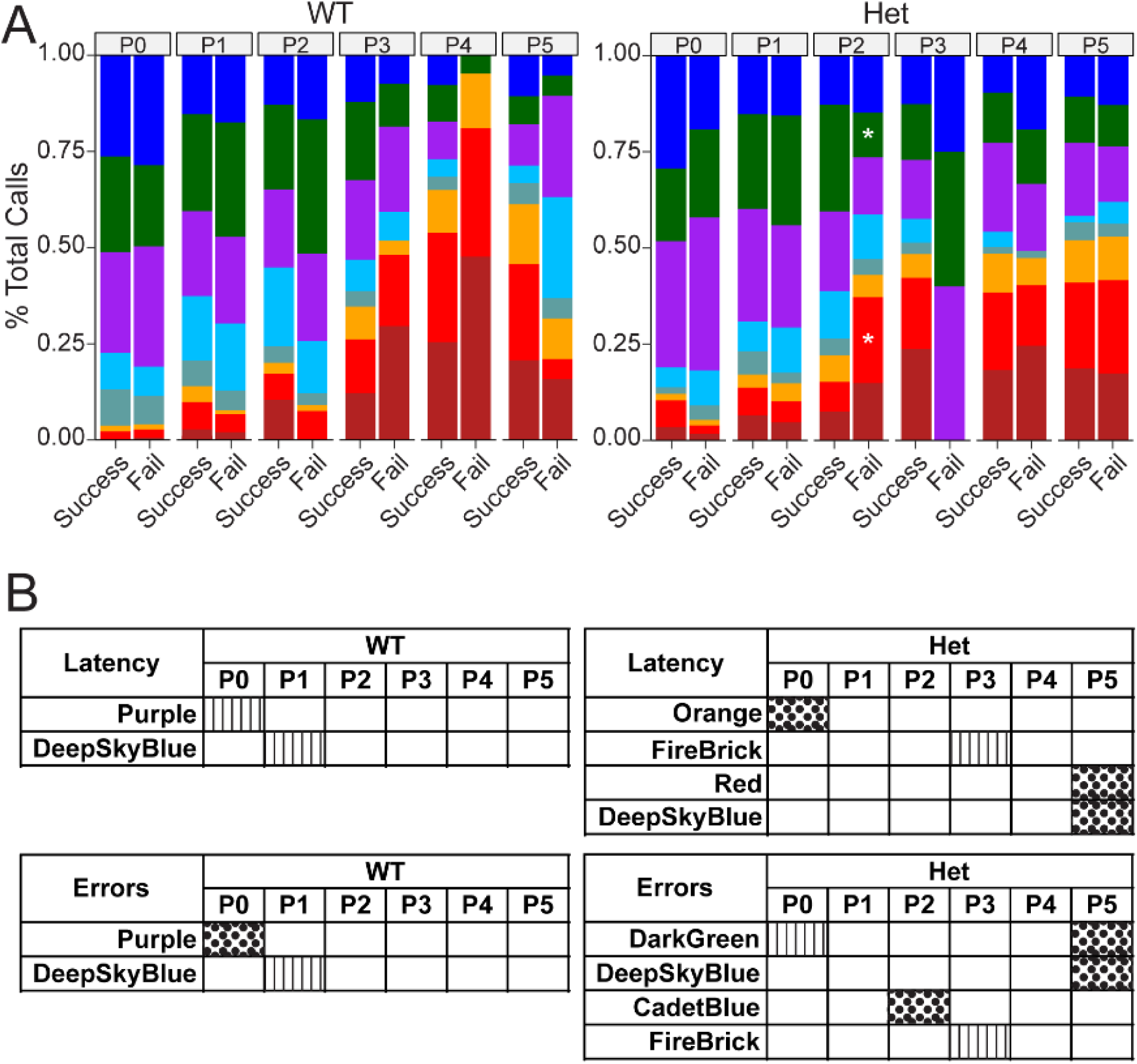
Pup call utilization shows limited correlation with behavioral outcome and efficiency. **A)** Bar plots depicting each cluster as a percentage of calls across each day between success and failure in WT (left) and Het (right). Statistical testing ability is limited due to limited number of calls, especially in WT fails, where fishers test fails to show significance in any day when corrected. *Fisher’s test: *p<0.05.* **B)** Statistical correlational analyses within genotypes between latency and errors and call cluster (linear regression). No cluster showed consistent significant correlation across days. Patterned boxes indicate significant correlation: vertical lines = significant negative correlations, and dotted boxes = significant positive correlations (*p<0.05*).

To characterize the sequential structure of pup USV calls (Figure 5-7), transition probabilities were calculated between all eight DeepSqueak clusters within each session. For each pair of consecutive calls within a session, a transition was counted if the interval between the end of the first call and the beginning of the second was 5 seconds or less; call pairs exceeding this window were treated as independent events. As calls are generally less than 1 second apart, the 5-second window was used as a filter to prevent any unrelated call events from being considered transitions. Raw transition counts were aggregated across sessions within each condition of interest (genotype, phase, day, and retrieval outcome) and converted to transition probabilities by dividing each row’s counts by the row total, yielding a matrix in which each entry reflects the probability of transitioning from one cluster to another given the source cluster. To correct for differences in cluster prevalence across conditions, transition probabilities were normalized to a null expectation defined as the overall frequency of the target cluster within that condition. Enrichment was calculated as the ratio of the observed transition probability to this null expectation, such that values greater than 1.0 indicate that transitions to a given target cluster occur more frequently than would be expected by chance given its prevalence, and values less than 1.0 indicate suppression. Enrichment values were capped at 6.0 to limit the influence of extreme ratios arising from very low-prevalence clusters. Panels with fewer than 5 calls in a given condition were excluded from visualization. For the grouped day analyses, calls from P1 through P3 were pooled to represent the mid postnatal period, and calls from P4 through P5 were pooled to represent the late postnatal period, with P0 analyzed separately as the early period.

**Figure 5.**
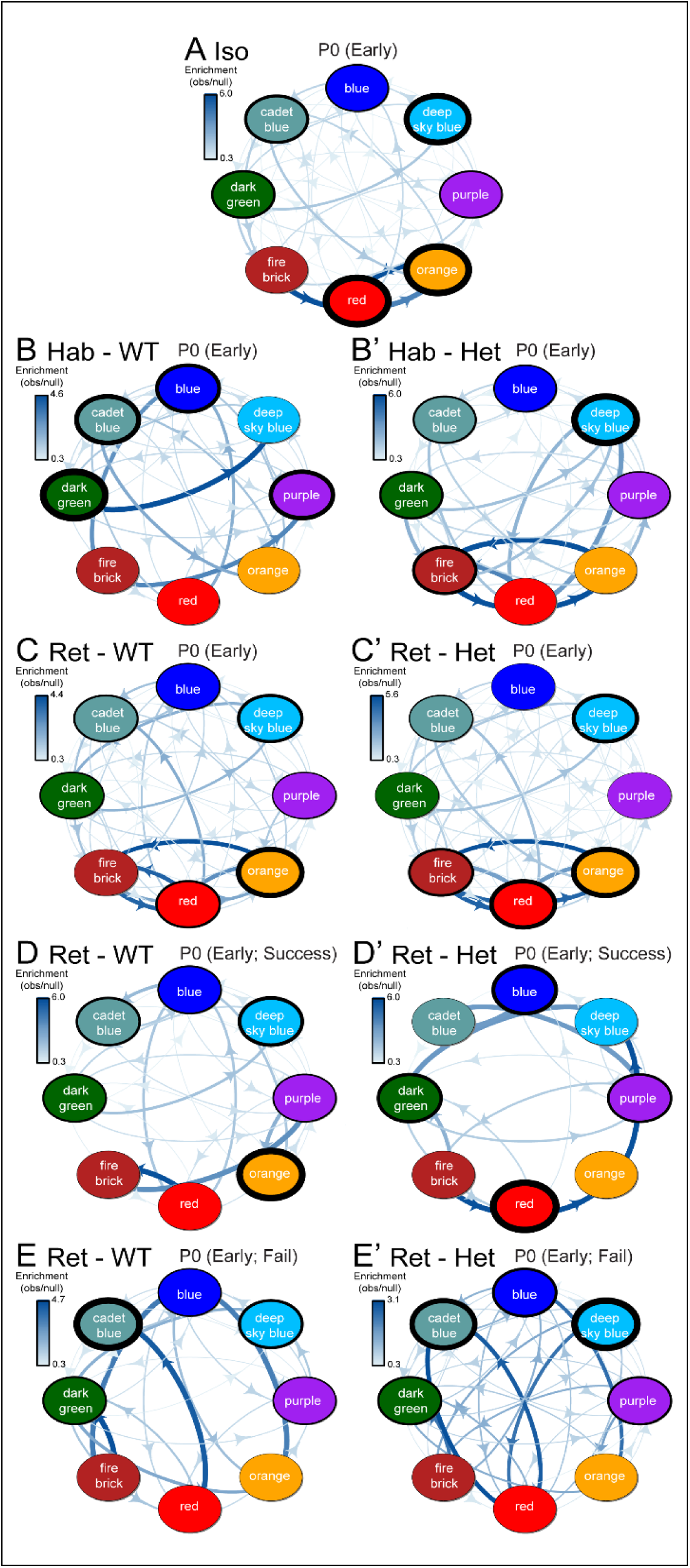
Call cluster transition enrichment at P0 (Early). Transition enrichment networks depicting the probability of sequential call cluster transitions during pup retrieval behavior at postnatal day 0 (P0), organized by behavioral phase and surrogate genotype. Arrow thickness and color intensity reflect enrichment above the null expectation (observed/null); arrows below the 0.3 enrichment threshold are not shown. Node outline thickness reflects self-enrichment (the degree to which a cluster transitions back to itself above chance). **(A)** Isolation phase (all pups, no surrogate present). **(B, B’)** Habituation phase with WT (B) and Het (B’) surrogates. **(C, C’)** Full 5-min retrieval phase with WT (C) and Het (C’) surrogates. **(D, D’)** Successful retrieval bouts with WT (D) and Het (D’) surrogates. **(E, E’)** Failed retrieval bouts with WT (E) and Het (E’) surrogates.

## Results

### Phase and day-specific dynamics in pup call rate during pup retrieval behavior

As published data typically focuses on isolation calls after P5, here, we collected and analyzed ∼59,000 individual USVs across the first 6 postnatal days and 3 phases of alloparental behavior (habituation, retrieval and pup isolation) using a YOLOv2 detection architecture (Redmon & Farhadi, 2018) built into DeepSqueak (Coffey et al., 2019). We found that from P3-P5 the pups emitted a significantly increased number of calls in isolation, which increased over the 6 days (Figure 1A). However, the number of calls remained consistent over days during habituation and retrieval phases, with both genotypes. (Figure 1A) We then performed linear regression analysis to determine the relationship between call rate, phases and days (Figure 1B, left). In isolation phase, call rates increased steadily from P0-P5 (R = 0.77, p = 3.3×10⁻⁸). However, call rate significantly decreased across days in the habituation phase (R = −0.27, p = 0.028), and remained stable in retrieval phase (R = -0. 0.069, P = 0.57). When call rates were separated based on adult genotype during retrieval, we found a similar yet insignificant trend (Figure 1B, right). The linear regression lines also had overlapping confidence intervals, indicating pup vocalization rate remained similar across P0-P5 regardless of surrogate genotype. Taken together, the stability of call rate over days and the lack of difference between surrogate genotypes indicates that pup retrieval performance differences between WT and Het are unlikely to be the result of pup call rate. Additionally, these results indicate that pups can produce USVs at different rates during contexts, through the first postnatal week.

USV emittance and a variety of behavioral responses of the surrogates occur over seconds during the dynamic pup retrieval behavior (Stevenson et al., 2021). Thus, we plotted the binned call rate across all phases and day (Figure 1C-D). We found striking temporal profiles at P0 vs. P5, with gradual changes over the other days (Supplemental figure 1). Specifically, during retrieval on P0, pup call rates were elevated early in the session when both WT and Het surrogates were present, and declined gradually over the session, likely indicating progression in retrieval. Temporal patterns of the pup calls were similar in isolation and habituation. By P5, temporal call rate profiles had diverged substantially: (1) isolation phase calls were highly elevated throughout, compared to all other phases; and (2) profile of pup calls during Het retrieval was also elevated, lacking the decline after 30 seconds with WT retrieval. This is likely a reflection of inefficient pup retrieval by Het on P5. Together, these results identify novel phase- and day-specific changes in pup call rate during pup retrieval behavior within the first week of life.

### DeepSqueak clusters are distinct in call metrics, separating into ‘cool’ and ‘warm’**categories**

The developmental changes in call rate over the first six days prompted us to question if the types of calls emitted changed over days. To answer this question, we used DeepSqueak to cluster and separate the ∼59000 calls in this data set utilizing Deepsqueak’s contour invariant unsupervised clustering by variational autoencoders (VAE); this approach yielded 8 distinct clusters. To interpret latent differences in clusters, descriptive acoustic metrics were remapped and visualized as radar plots (Fig 2A). These metrics revealed two major groups, ‘warm’ and ‘cool’ clusters, as denoted by the colors chosen to represent each cluster.

The cool group (blue, deep sky blue, cadet blue, dark green, purple) displays lower principal frequency and longer call lengths. The warm group (red, orange, fire brick) shows high principal frequencies, frequencies ranging between 85-110k kHz, short call lengths, and minor call slopes and sinuosity. Furthermore, the cluster characteristics remain predominantly similar (except for deep sky blue which shows higher sinuosity when Het are present), regardless of surrogate genotype. To determine if there were day-specific changes in the clusters, we plotted the distance from the center as a measurement of each call’s eigenvalue distance from the most “central” or average call of the cluster. Each cluster’s median distance from the center shows its distinctness. Most clusters’ medians are around 2.5, indicating that the clusters are similar around their own centroids (Suppl. Figure 2). Clusters’ distance does not change significantly across the course of development, except for purple. After correcting for multiple comparisons using Benjamini Hochberg test, cluster characteristics do not vary differentially between phases (Table 1). The most representative call for each cluster was extracted from DeepSqueak and plotted for call length and frequency (Figure 2B). The separation of warm and cool clusters is distinct, with warm calls displaying typically short duration with high frequency whereas the cool calls are longer with lower frequency. When call length and slope are plotted (Figure 2C), the shapes of the representative calls are evident with most clusters showing downward slopes (except red and purple, which are flat). Together, these results introduce the 8 clusters and their characteristics throughout development, phases and genotypes.

**Table 1:** Two-sided Wilcoxon rank-sum tests compared phase (Hab, habituation; Ret, Retrieval, 5 min retrieval; Iso, isolation) for seven acoustic parameters within each cluster. “Significant before correction” lists parameters with uncorrected p < 0.05. p-values were then Bonferroni-corrected across all 168 tests; no parameter reached significance at adjusted alpha = 0.05 in any comparison (ns).

| **Cluster** | **Comparison** | **Significant before correction** | **Significant after correction** |
|---|---|---|---|
| **Blue** | Hab vs Ret | Call Length, High Freq, Delta Freq, Sinuosity | *NS* |
|  | Hab vs Iso | High Freq | *NS* |
|  | Ret vs Iso | *NS* | *NS* |
| **Dark green** | Hab vs Ret | Call Length, High Freq, Delta Freq | *NS* |
|  | Hab vs Iso | *NS* | *NS* |
|  | Ret vs Iso | *NS* | *NS* |
| **Purple** | Hab vs Ret | Call Length, High Freq, Delta Freq, Sinuosity | *NS* |
|  | Hab vs Iso | High Freq, Delta Freq | *NS* |
|  | Ret vs Iso | *NS* | *NS* |
| **Orange** | Hab vs Ret | Call Length, Principal Frequency, High Freq, Delta Freq | *NS* |
|  | Hab vs Iso | Call Length, High Freq, Delta Freq | *NS* |
|  | Ret vs Iso | *NS* | *NS* |
| **Red** | Hab vs Ret | Call Length, Principal Frequency, High Freq, Delta Freq, Sinuosity | *NS* |
|  | Hab vs Iso | Call Length, High Freq, Delta Freq | *NS* |
|  | Ret vs Iso | *NS* | *NS* |
| **Deep sky blue** | Hab vs Ret | Call Length | *NS* |
|  | Hab vs Iso | *NS* | *NS* |
|  | Ret vs Iso | Call Length | *NS* |
| **Cadet blue** | Hab vs Ret | *NS* | *NS* |
|  | Hab vs Iso | *NS* | *NS* |
|  | Ret vs Iso | Call Length | *NS* |
| **Fire brick** | Hab vs Ret | High Freq, Delta Freq | *NS* |
|  | Hab vs Iso | Call Length, High Freq, Delta Freq | *NS* |
|  | Ret vs Iso | Call Length | *NS* |

### Call clusters significantly change across phases of behavior and days

Next, we asked if the individual call clusters were more prevalent in the different phases of pup retrieval behavior. Specifically, to determine if certain call clusters play an important role in the retrieval behavior, we chose to investigate call cluster distribution across days in each phase of the pup retrieval behavior. In habituation, we find 5/8 clusters showing no significant changes over days (Figure 3A, top). Cool clusters (blue, purple and cadet blue) show significant decreases over days). In retrieval, we find an overall increase in warm group (3/3) and a decrease in cool group’s prevalence over days (except for deep sky blue, 4/5) (Figure 3A, bottom). We found that, in the isolation phase, number of pup calls was influenced by development, with a significant increase in warm group’s prevalence (3/3), and modest increase in cool group’s prevalence (3/5, only in purple, blue and deep sky blue) (Figure 3A, middle). Together, these results suggest that though pup call development over days has a strong influence, behavioral context is key for modulating pup call counts even in this first week of life.

When the WT surrogate was present in the home cage with the pups, either in habituation or retrieval, we found a significant decrease in cool calls (4/5 clusters on most days) and day-specific increases in warm clusters (red on P2, P4. P5; orange on P2, P4, P5) (Figure 3B, left). When the Het surrogate is present with the pups, we found a pattern similar to the WT, though to a lesser extent (Figure 3B, right). These average patterns are well illustrated in sequence plots of a representative example across days and phases (Figure 3C, top). During isolation, the prevalence of the cool group in early days and dominance of the warm group in later days is clear. Even with an adult of either genotype present in the cage and the pups huddled in the nest together (in habituation phase, Figure 3C, bottom), pups maintain the prevalence of early cool clusters and later warm clusters. During the retrieval phase, pups continue to call most days, following the same pattern (Figure 3C, middle). Together, these results suggest that the pup calls’ prevalence changes in a context-dependent manner, within the constraints of the developmental profile, regardless of the genotype of the surrogate.

### Call clusters are not consistently associated with retrieval outcome or efficiency

Next, we questioned if any cluster prevalence changed with success or failure in retrieval behavior (Figure 4A). Retrieval events were scored as success or fail events based on whether or not the event ended in surrogate returning the pup to nest. Events were timestamped from initial interaction with the pup (whisker length away) until pup was successfully retrieved or dropped in the case of failures. All calls occurring during the timestamped event were labeled as either ‘success’ or ‘fail’ depending on outcome of event.

With WT surrogates, the call clusters were similar during successful and failed retrieval events (Figure 4, left). With Het surrogates, we noted statistically significant changes in red and dark green clusters during failed retrievals, only on P2 (Figure 4, right; p = 0.023). When the calls during successful or failed retrievals were normalized to account for individual day prevalences (Table 2), additional patterns emerged that were not apparent from raw call counts alone. With WT surrogates, both warm clusters (orange, red) and cadet blue showed the strongest enrichment during successful retrievals at P0 and P1, reflecting a pattern in which rare call types are disproportionately associated with successful retrieval events in early days of retrieval. Enrichment for these clusters declined over days, while other cool clusters showed increasing enrichment by P5. Calls associated with failed retrievals with WT surrogates were minimal (n = 1-4 calls on most days), limiting interpretation. With Het surrogates, regardless of outcome, cluster enrichment was more evenly distributed across all clusters and generally lower, with fire brick, orange, and cadet blue showing the most consistent enrichment across days.

**Table 2.** Enrichment of clusters during pup retrieval bouts, by outcome, genotype, and postnatal day. Values represent the relative enrichment (observed probability ÷ null expectation based on cluster prevalence) for each of the eight clusters across postnatal days P0–P5. Values greater than 1.0 indicate that a given cluster is called at a rate exceeding chance (bolded, underlined); Dashes indicate insufficient calls for a reliable estimate. *n* = number of retrieval bouts contributing to that day’s estimate.

| WT | Success |  |  |  |  |  | Fail |  |  |  |  |  |
|---|---|---|---|---|---|---|---|---|---|---|---|---|
| **Cluster** | **P0** | **P1** | **P2** | **P3** | **P4** | **P5** | **P0** | **P1** | **P2** | **P3** | **P4** | **P5** |
| *n =* | *16* | *23* | *19* | *12* | *25* | *18* | *4* | *7* | *4* | *1* | *1* | *2* |
| **Blue** | 0.71 | **1.98** | 0.82 | **2.74** | 0.52 | **2.6** | **1.75** | **1.64** | **1.5** | — | — | **9.5** |
| **Dark green** | 0.76 | 0.17 | 0.72 | 0.41 | 0.84 | **1.52** | **1.19** | 0.48 | 0.72 | — | — | **9.5** |
| **Purple** | 0.71 | 0.59 | 0.52 | 0.4 | **1.22** | **1.04** | 0.8 | 0.63 | — | — | — | — |
| **Deep sky blue** | 0.66 | 0.26 | **1.28** | **1.03** | 0.89 | **1.21** | — | — | **1.83** | — | — | — |
| **Cadet blue** | **2.65** | **3.23** | **1.23** | **2.06** | **1.16** | — | — | **2.79** | — | — | — | — |
| **Orange** | **4.2** | — | **3.68** | 0.97 | **1.08** | 0.71 | — | — | — | — | — | — |
| **Red** | **3.15** | **2.46** | **2.36** | **1.19** | 0.7 | 0.67 | — | **6.19** | — | — | **3** | — |
| **Fire brick** | — | **3.15** | 0.5 | 0.69 | **1.41** | 0.8 | — | — | — | **3.38** | — | — |
| **Het** | **Success** |  |  |  |  |  | **Fail** |  |  |  |  |  |
| **Cluster** | **P0** | **P1** | **P2** | **P3** | **P4** | **P5** | **P0** | **P1** | **P2** | **P3** | **P4** | **P5** |
| *n =* | *6* | *23* | *23* | *15* | *21* | *16* | *29* | *25* | *9* | *1* | *4* | *19* |
| **Blue** | **1.13** | **1.25** | **1** | 0 | **1.61** | **1.32** | **1.3** | **1.21** | 0.96 | 0 | **1.52** | **1.31** |
| **Dark green** | 0.25 | 0.91 | **1.06** | 0 | 0.8 | **1.24** | 0.27 | **1.05** | **1.16** | 0 | 0.74 | **1.21** |
| **Purple** | **1.17** | 0.59 | **1.3** | 0.77 | 0.53 | 0.73 | **1.26** | 0.52 | **1.46** | 0 | 0.49 | 0.71 |
| **Deep sky blue** | **1.8** | 0.58 | 0.45 | 0.97 | **1.26** | 0 | **1.95** | 0.49 | 0.38 | 0 | 0.93 | 0 |
| **Cadet blue** | 0.93 | **1.51** | 0.86 | **2.53** | 0 | **1.26** | **1.04** | **1.37** | 0.54 | — | 0 | **1.15** |
| **Orange** | 0 | **3.08** | **1.51** | **2.58** | 0.34 | 0.87 | 0 | **2.79** | **1.79** | 0 | 0.41 | 0.97 |
| **Red** | 0 | 0.57 | 0.68 | **1.4** | **1.11** | 0.95 | 0 | 0.72 | 0.56 | 0 | **1.16** | 0.93 |
| **Fire brick** | 0 | **1.75** | **1.4** | **1.29** | **1.61** | **1.43** | 0 | **1.59** | **1.5** | 0 | **1.42** | **1.45** |

Correlational analysis between normalized clusters and surrogate retrieval efficiency revealed only day-specific associations (Figure 4B). With WT retrieval metrics of latency and errors, cool clusters purple and deep sky blue had significant correlations with only the first 2 days of retrieval metrics. Het showed a different pattern, with latency metrics significantly and positively correlated with orange, red and deep sky blue on different days while negatively correlated with fire brick only on P3. Error metrics had significant correlations with cool clusters (3/5) and with fire brick on P3. Together, these results indicate that call cluster identity is not consistently associated with behavioral outcome or efficiency with either surrogate genotype.

### Call sequences show context-dependent structure

Since individual call clusters did not consistently discern phases or efficient retrieval, we asked if the sequence of calls, measured by transition probability, would reveal distinct patterns in calls and behavioral outcome (Figure 5-7). At P0, during habituation with WT surrogate (Figure 5B), pup USVs had strong transition probability between deep sky blue and dark green clusters, and self-transitions in dark green, cadet blue, blue and purple clusters. With Het present (Figure 5B’), pup USVs showed strong transition probability in the warm clusters, and strong self-transitions in deep sky blue cluster. A combination of these patterns was found in the isolation phase (Figure 5A), with strong transitions between warm clusters, and strong self-transitions in deep sky blue, orange and red clusters. This pattern carried over to the retrieval phase with both WT and Het (Figure 5C, C’) and diverged in successful and failed retrievals (Figure 5D-E’). Though consistent patterns did not emerge in the successful transition probability plots between WT and Het (Figure 5D, D’), the transition probability lines crossed cool and warm clusters in failure events for both genotypes (Figure 5E, E’). A caveat for the WT failed data is that their transition probability lines were generated from only 1-4 USVs in these clusters, as WT surrogates had minimal failed interactions. Together, these plots suggest that even on P0, phase-specific transition probability in call clusters occurs, with emphasis on warm cluster calls in isolation and retrieval, regardless of genotype.

**Figure 6.**
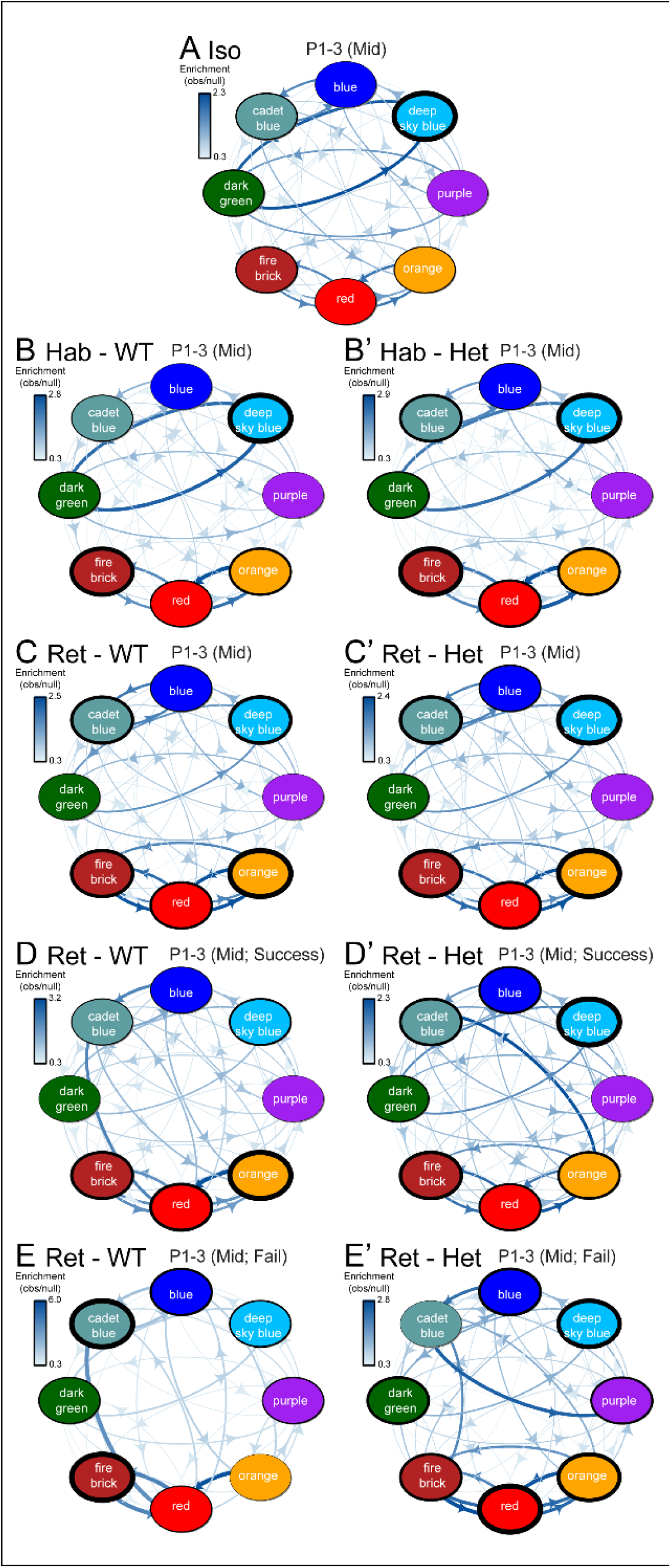
Call cluster transition enrichment at P1–3 (Mid). Transition enrichment networks depicting the probability of sequential call cluster transitions during pup retrieval behavior across postnatal days P1–P3, pooled to represent the mid postnatal period. Layout and encoding are identical to Figure 6. **(A)** Isolation phase. **(B, B’)** Habituation phase with WT (B) and Het (B’) surrogates. **(C, C’)** Full 5-min retrieval phase with WT (C) and Het (C’) surrogates. **(D, D’)** Successful retrieval bouts with WT (D) and Het (D’) surrogates. **(E, E’)** Failed retrieval bouts with WT (E) and Het (E’) surrogates.

**Figure 7.**
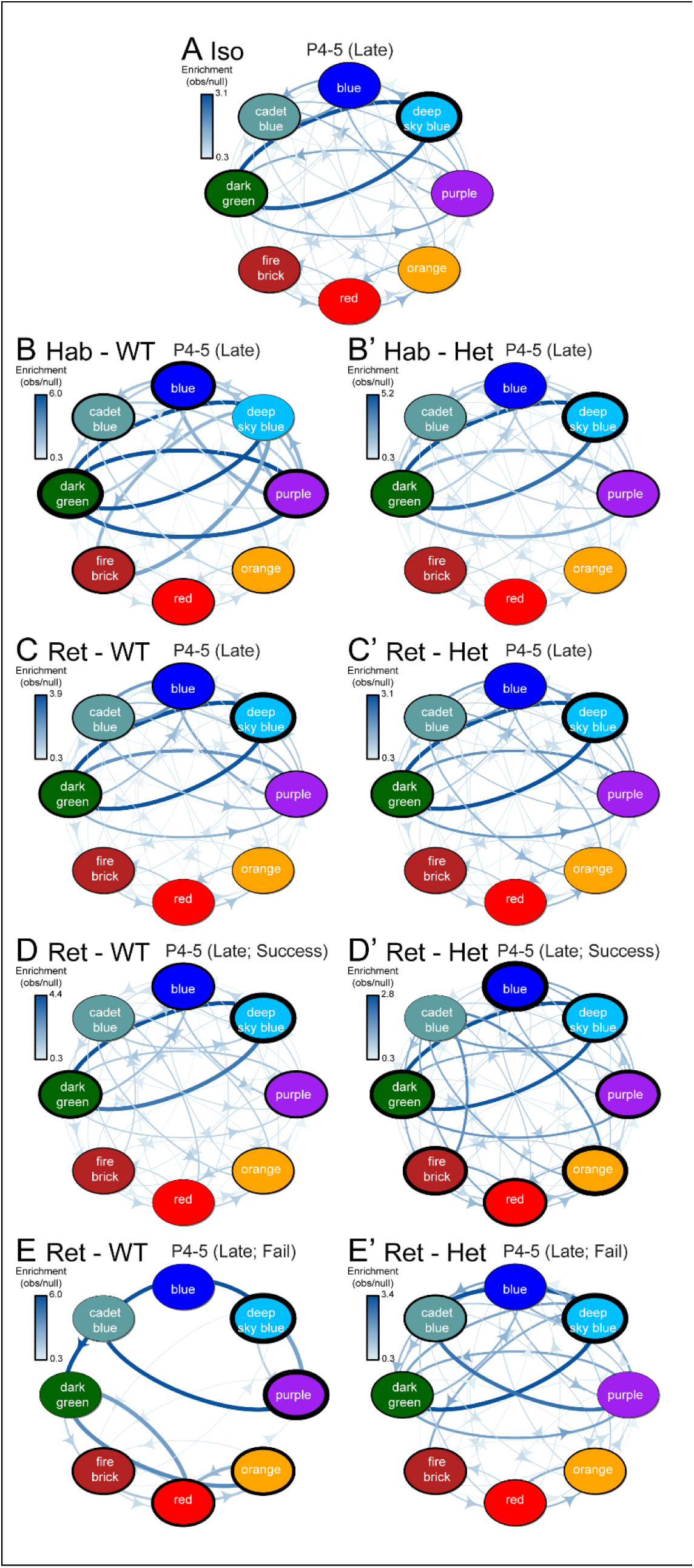
Call cluster transition enrichment at P4–5 (Late). Transition enrichment networks depicting the probability of sequential call cluster transitions during pup retrieval behavior across postnatal days P4–P5, pooled to represent the late postnatal period. Layout and encoding are identical to Figure 6. **(A)** Isolation phase. **(B, B’)** Habituation phase with WT (B) and Het (B’) surrogates. **(C, C’)** Full 5-min retrieval phase with WT (C) and Het (C’) surrogates. **(D, D’)** Successful retrieval bouts with WT (D) and Het (D’) surrogates. **(E, E’)** Failed retrieval bouts with WT (E) and Het (E’) surrogates. All pups are wild-type; WT and Het refer to the genotype of the retrieving dam.

Next, we examined the transition probabilities of call clusters during the middle stages of maternal experience. We combined days P1-P3 data as they had similar patterns (Figure 6, ’mid’ days). In mid days, the transition probabilities in the call clusters across phases became more apparent and consistent. Deep sky blue and dark green transitions, and warm color transitions dominated habituation and isolation phases (Figure 6A-B’). Consistent warm color transitions and deep sky blue self-transitions were prevalent in Retrieval and Success (Figure 6C-D’). Failure events still showed strong cool – warm cluster transitions, typically not seen in any other phase. Despite this unique pattern, these patterns are unlikely to contribute to the surrogates’ performance in the retrieval behavior, consistent with previous findings in this study.

Lastly, we examined the transition probabilities of call clusters during the late stages of maternal experience. We combined P4-P5 days (Figure 7, ‘late’ days) and showed that the call cluster transition probabilities were stabilized but different from the earlier patterns. The warm cluster transitions were lost across all phases (Figure 7A-E’). The transitions between deep sky blue and dark green remained strong in all phases, with deep sky blue self-transition dominating. The cool-warm transition lines seen in failure events above were minimal, mainly in WT Fail, which was a rare event (Figure 7E). Moreover, transition patterns were similar with WT and Het during habituation and retrieval (Figure 7B-C’).

Together, these results suggest that the pup calls develop their patterns of call sequences over the first week, with context-dependent transitions even on the first day. Given that cool cluster calls dominate in early days, and warm calls dominate in later days (Fig. 3), this opposite call sequence pattern (with warm transition probability lines in early days becoming minimal in later days), though minimal in numbers, could constitute a “signal” within the cacophony of “noise” calls throughout the days.

## Discussion

Ultrasonic vocalizations represent a foundational model for understanding social communication development, with direct relevance to neurodevelopmental disorders affecting language and social behavior. Despite extensive characterization of pup calls across numerous genetic models, there is still no consensus on pup call types, and how they impact maternal care. Previous classification efforts have typically sampled a few thousand calls in a single context, typically in isolation, and at individual days, leaving the developmental and behavioral contextual structure of the pup call repertoire unresolved. Here, we addressed this question by characterizing over 59,000 pup calls across P0-P5 during pup retrieval paradigm. This is one of the largest reported data sets for a consecutive postnatal study, reflecting daily sampling across behavioral context. Using unsupervised, VAE-based clustering, we identified eight call types whose relative prevalence shifted systematically with both postnatal age and behavioral context, providing a detailed developmental map of the early pup call repertoire. Our findings support a model in which USVs serve as general arousal signals, that are then combined with other sensory inputs to elicit maternal care. Furthermore, by examining pup calling in the context of impaired maternal care from *Mecp2*-heterozygous females, we demonstrate that sensory processing capabilities in the adult Het surrogate likely, rather than pup vocal adjustments, determine retrieval outcomes.

### Pup USVs exhibit dynamic developmental trajectory

Audio data are highly complex, and typical descriptive features utilized by the field are often highly correlated. Variational Autoencoders address this by learning latent features directly from the data, rather than relying on hand-selected, correlated descriptive features. Auto-encoder based classification outperforms contour characteristic-based clustering (Goffinet & Pearson, 2021). The VAE-based clustering captured the developmental trajectory of the pup vocal repertoire, identified eight distinct clusters that shifted systematically as pups matured. Low-frequency, high-slope calls (cool clusters) dominated early days, while high-frequency, short-duration calls (warm clusters) increased progressively and comprised the majority by P4-P5. This shift likely reflects maturational changes in mouse larynx structure, respiratory control and neural coordination (Grimsley et al., 2011; Håkansson et al., 2022; Heckman et al., 2016; Riede et al., 2020). Importantly, this pattern was consistent across all behavioral contexts and occurred identically regardless of surrogate genotype, suggesting maturation of vocal systems rather than experience-dependent plasticity drives this transition. This is evident in the presence and increasing prevalence of higher frequency calls over the six days. The eight-cluster solution represents a methodological advance over previous categorization efforts yielding 6 to over 100 categories depending on feature selection (Dou et al., 2018; Elwood & Keeling, 1982; Portfors, 2007; Sales & Smith, 1978). Our clusters showed strong stability across validation approaches and mapped interpretably onto established acoustic parameters.

### Call sequences show context-dependent structure

Although individual call clusters did not discriminate with behavioral phase or retrieval outcome, the sequence in which clusters were produced showed clear phase- and development-specific structure throughout the six days. Habituation and isolation were characterized by transitions linking cool and warm clusters, whereas retrieval showed more stable self-transitions within a cluster type. Differential calling in these phases suggests that pups are capable of distinguishing contexts in this first week of life. Pup USV generation is thought to result from lack of body temperature self-regulation (Okon, 19070). Due to the call cluster differences observed in retrieval and isolation phases, our results suggest that pups may be able to distinguish when an adult caretaker is present with them, potentially via vibrations or in response to tactile interaction from the surrogates, as pup ear holes do not open until P4. Furthermore, by P4-P5, the warm-cluster transitions were largely lost and cool-cluster self-transitions dominated. The clearest candidate for the behaviorally relevant signature was a cool-to-warm transition specific to failed retrieval attempts in the early and middle postnatal days, in both genotypes, which faded by P4-P5 as warm cluster calls became prevalent overall (Fig. 3). This is a dissociation between raw call-type prevalence and sequence structure. As these transitions were from a few calls, particularly with WT failures, we propose this as a hypothesis-generating observation. This type of sequence-level analysis has been established for adult vocalizations, where courtship song syntax shifts with social context and predictive call sequences are linked to auditory cortex tuning (Agarwalla et al., 2023; Chabout et al., 2015; D’amato, 1991; Egnor & Seagraves, 2016; Hanson & Hurley, 2012; Holy & Guo, 2005). To our knowledge, this specific approach, of linking call-type transition probability to postnatal development and pup retrieval behavioral outcome, has not been applied to pup calls in such an ethologically relevant manner. This analysis adds a temporal, sequence-level layer to the mouse pup call-type atlas.

### No distinct call signature triggers efficient retrieval behavior

Our central finding is that no specific call cluster consistently predicted or facilitated maternal retrieval behavior. This contrasts with adult rodent vocalizations, where specific call types are associated with distinct social contexts, suggesting categorical specificity emerges later as production and perception systems mature (Agarwalla et al., 2023; D’amato, 1991; Hanson & Hurley, 2012; Portfors, 2007). Adult rats have been identified to have a variety of context specific call types within the 50Hz band (Wright et al, 2010; Wöhr et al., 2008b). There is some evidence supporting context specific call types in mice as well, specifically in adult males, in social contexts, courtships, and behavioral state (Chabout et al., 2015; Egnor & Seagraves, 2016). Despite clear increases in overall call rate during isolation phase over the first six days of life, no call cluster is uniquely expressed in a phase-specific or genotype-specific manner. However, the composition of the call clusters changes dynamically over days tied to one phase. Correlation analyses revealed a limited relationship between any clusters and retrieval latency. This pattern is held when restricting analysis to successful versus failed attempts. Rather than serving as unique, categorized signals specifying maternal actions, pup calls appear to function primarily as general alert signals. This interpretation aligns with evidence that maternal retrieval depends on multisensory integration. Olfactory cues are also necessary for retrieval, and pup USVs potentiate maternal responsiveness to olfactory stimuli rather than eliciting behavior independently (McRae et al., 2023; Okabe et al., 2013; Shair et al., 1998; Smotherman et al., 1974). USVs direct maternal attention and approach, while other sensory inputs, such as olfactory and tactile cues, provide specificity needed to guide retrieval actions (McRae et al., 2023; Smotherman et al., 1974). Considering previous work in our lab and others showing processing deficits in auditory, tactile, and olfactory cues in Het, impacted multisensory processing and integration are likely to be the cause of their reduced ability to retrieve (Brima et al., 2019; Lau et al., 2020a; Lau et al., 2020b; Mykins et al., 2023; Rupert & Shea, 2022; Stevenson et al., 2021).

We tested whether pups adjust calling in response to impaired maternal care from *Mecp2*-heterozygous surrogates. Previous work demonstrated that Het surrogates show dramatically prolonged retrieval latencies, increased errors, and frequent failures, compared to WT surrogates (Krishnan et al., 2017; Mykins et al., 2024; Stevenson et al., 2021). However, across all six testing days and analytical breakdowns, pups showed no systematic differences in cluster distribution when interacting with WT vs Het surrogates. While individual clusters showed significant differences on specific days, these effects were inconsistent, and do not typically withstand multiple corrections testing. Critically, this demonstrates that impaired maternal behavior cannot be attributed to inadequate pup signals, as Het surrogates receive nearly identical vocal input yet fail to respond appropriately.

The finding that pups do not alter calling combined with pronounced Het maternal deficits provides additional evidence that heterozygous MeCP2 expression disrupts maternal sensory processing rather than pup signal production. This aligns with our previous neurophysiological findings showing surrogate Het females fail to discriminate pup calls from pure tones in primary auditory cortex (Krishnan et al., 2017). While WT females discriminately showed enhanced excitatory and reduced inhibitory responses to calls over pure tones, Het females showed altered responses to both excitatory and inhibitory responses, with a significant increase in inhibitory activity, to both calls and pure tones (Krishnan et al., 2017; Lau et al., 2020a), indicating a failure to discriminate socially relevant acoustic signals. Pups provide identical acoustic input to WT and Het surrogates, yet Het surrogates show dramatically impaired retrieval efficiency, highlighting the deficit within maternal sensory processing and sensorimotor integration systems.

### Implications for USVs in the social communication field

These results refine our understanding of USV function during early development and highlight the importance of coupling acoustic characterization with behavioral validation. A common approach has been characterizing altered USVs in genetic models and interpreting these alterations as communication deficits. Our findings suggest this interpretation requires caution. Changes in acoustic parameters or call type distributions may not impact social communication outcomes if those features are not functionally relevant to the receiving individual. In contrast, our Het model demonstrates a true communication deficit: not because pups call abnormally, but because Het females cannot appropriately process and respond to normal pup calls. This highlights the necessity of examining both signal production and reception when characterizing social communication phenotypes.

### Limitations

A major limitation of our analysis is our recording setup, which does not allow for direct identification of single pup calls emitted. All microphones pick up calls from pups in any position in the cage. Therefore, it limits our ability to do certain analysis such as temporal dynamics within calling bouts or precise timing relative to maternal behaviors, which may influence maternal responses. Future experiments could include a one pup retrieval paradigm to directly investigate individual pup calls impact on maternal retrieval. However, we used this scattered multiple-pups paradigm to reflect more naturalistic environment where an adult mouse would have to gather and take care of multiple pups at the same time. Future work utilizing sequence playback experiments will test whether neurons respond differentially in a development-specific manner.

## Author contributions

KK supervised the project, data analysis, designed experiments and contributed to writing of the manuscript. JE performed data analysis, visualizations and writing of the manuscript. JB performed data analysis and initial results. KJR, DC and BYBL contributed to research design, analysis and discussions. KK and BYBL also collected data.

## Acknowledgments

We would like to thank the following undergraduate and graduate students for their help with data collection and initial manual USV analysis. Brett Emery, Ashlee Tannehill, Matthew Everett, Dana Layo-Carris, Delaney Fisher, Sarah Roth, Parker Stevenson, Radha Awasthi, Raymond Senu, Benjamin Bridges and Hannah Lee. This work was supported through startup funds to KK, the National Institute of Mental Health of the National Institutes of Health under Award Number R15MH124042 (KK) and generous support for undergraduate research through the University of Tennessee.

---

**Supplemental figure 1:**
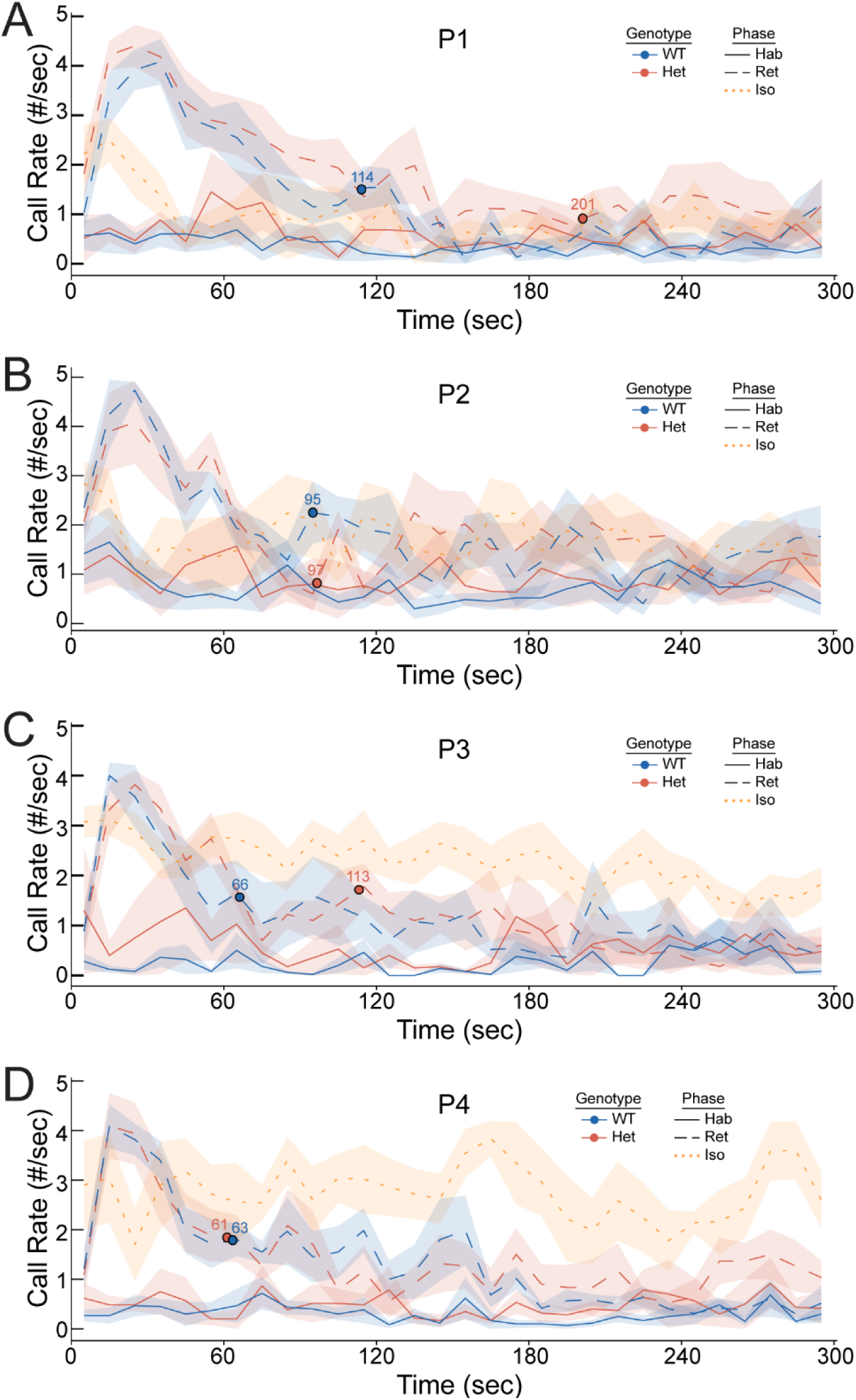
Line plots depicting calls binned per 10 seconds for WT (blue) and Het (red) in Hab (solid line) and Ret (dashed line) on P1 **(A)**, P2 **(B)**, P3 **(C)**, and P4 **(D)** for the full 300 seconds of the phase, with circles indicating average end of retrievals for each genotype. Iso (yellow, dotted) calls are binned similarly. Pup calls during WT retrieval showed decreases across days, while calls during Het retrieval showed variability across days. Iso calls increased in P3 and P4. Hab calls maintained low across days.

**Supplemental figure 2:**
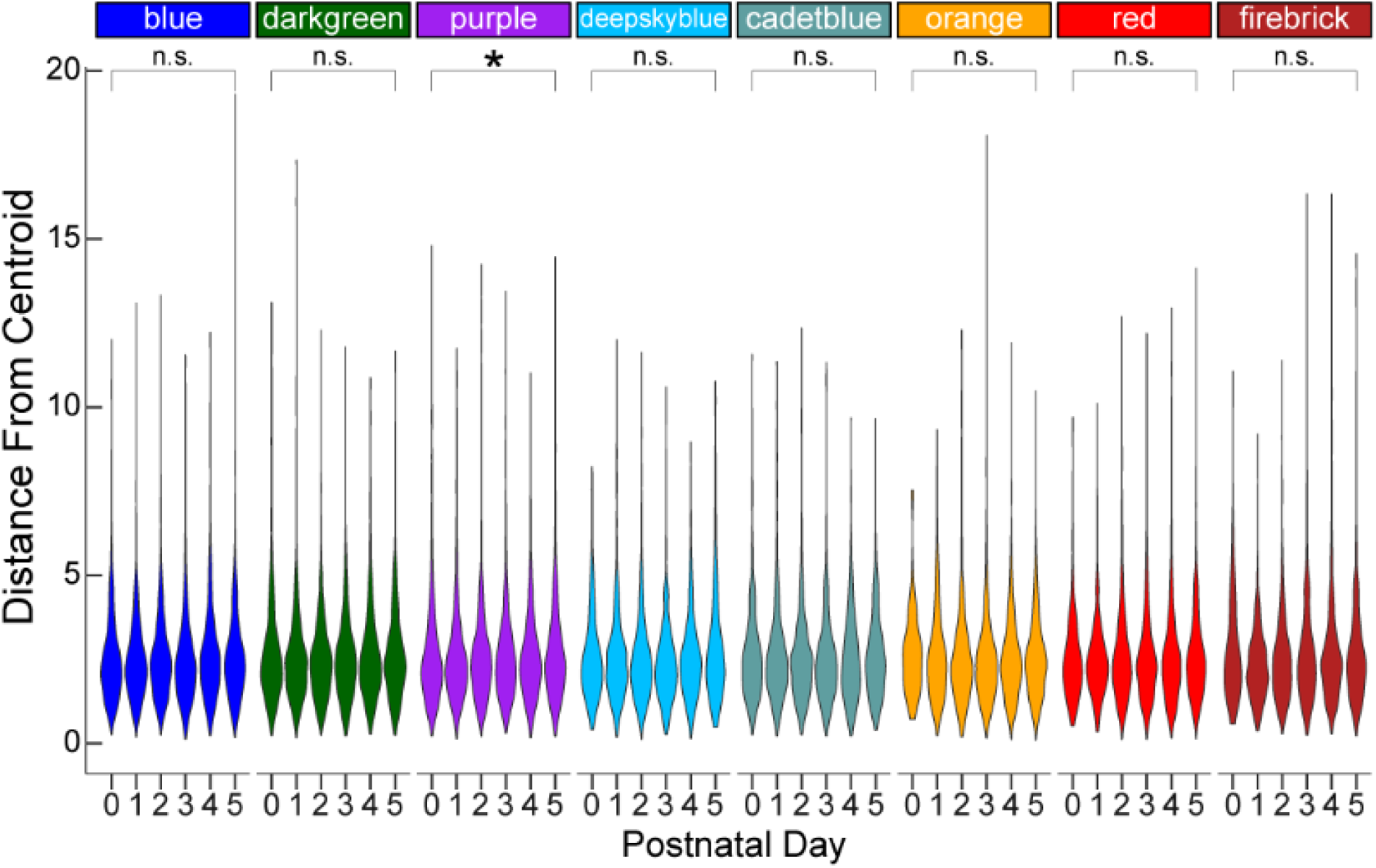
Violin plots indicating distribution of each call’s distance from center of its cluster across days of behavior. Purple cluster was the only cluster that showed significant increase in distance from center across time, indicating that calls remain consistently separate throughout the course of the behavior. Mean comparisons are between P0 and P5. *Mann Whitney, FDR corrected, * = p<.05*, n.s. = not significant

